# Towards robust evolutionary inference with integral projection models

**DOI:** 10.1101/064618

**Authors:** Maria João Janeiro, David W. Coltman, Marco Festa-Bianchet, Fanie Pelletier, Michael B. Morrissey

## Abstract

Integral projection models (IPMs) are extremely flexible tools for ecological and evolutionary inference. IPMs track the distribution of phenotype in populations through time, using functions describing phenotype-dependent development, inheritance, survival and fecundity. For evolutionary inference, two important features of any model are the ability to (i) characterize relationships among traits (including values of the same traits across ages) within individuals, and (ii) characterize similarity between individuals and their descendants. In IPM analyses, the former depends on regressions of observed trait values at each age on values at the previous age (development functions), and the latter on regressions of offspring values at birth on parent values as adults (inheritance functions). We show analytically that development functions, characterized this way, will typically underestimate covariances of trait values across ages, due to compounding of regression to the mean across projection steps. Similarly, we show that inheritance, characterized this way, is inconsistent with a modern understanding of inheritance, and underestimates the degree to which relatives are phenotypically similar. Additionally, we show that the use of a constant biometric inheritance function, particularly with a constant intercept, is incompatible with evolution. Consequently, current implementations of IPMs will predict little or no phenotypic evolution, purely as artifacts of their construction. We present alternative approaches to constructing development and inheritance functions, based on a quantitative genetic approach, and show analytically and through an empirical example on a population of bighorn sheep how they can potentially recover patterns that are critical to evolutionary inference.

## Introduction

Evolutionary and ecological dynamics converge at the scale of generation-to-generation change in populations (Pelletier et al., 2009; Coulson et al., 2010). When traits cause fitness variation, the distributions of those traits, weighted by fitness, necessarily changes within generations (Godfrey-Smith, 2007). If differences among individuals have a genetic basis, then genetic changes will be concomitant with phenotypic changes. Such genetic changes are the basis for the transmission of within-generation change due to selection, to genetic change between populations, i.e. evolution (Lewontin, 1970; Endler, 1986). The fundamental nature of this relationship between phenotypic change due to selection, and associated genetic and thus evolutionary change, has motivated the development of various expressions relating selection to genetic variation and evolution in quantitative terms (Lush, 1937; Robertson, 1966, 1968; Lande, 1979; Lande & Arnold, 1983; Morrissey, 2014, 2015). Important recent advances in population demography, particularly the introduction (Easterling et al., 2000) and popularization (e.g. Childs et al., 2003; Ellner & Rees, 2006; Coulson et al., 2010; Ozgul et al., 2010; Coulson, 2012; Merow et al., 2014) of integral projection models (IPMs), can potentially allow the construction of very flexible models of changes in phenotype, and of its associated demographic implications (Coulson et al., 2010).

IPMs are structured population models used to study the dynamic of populations when individuals’ vital rates (e.g. survival, growth, reproduction) depend on one or more continuous state variables (e.g. mass). In principle, these model structures track the distribution of individual values of the state variables through time. To achieve this, IPMs make population projections from regression models that define the underlying vital rates as a function of the state variables. Four core sets of functions for vital rates have been defined, termed fundamental functions or fundamental processes (Coulson et al., 2010): (i) survival, (ii) fertility, (iii) ontogenetic development of focal trait conditional on surviving (development functions), and (iv) distribution of offspring trait as a function of parental trait (inheritance functions). In principle, the inheritance functions allow IPMs to be used to make evolutionary inference, i.e. inference of evolutionary trajectories and parameters relevant to evolutionary processes, such as selection and genetic variation, particularly including the estimation of biometric heritabilities (Coulson et al., 2010; Schindler et al., 2013; Traill et al., 2014; Bassar et al., 2016). As discussed by Coulson et al. (2010), these four processes underlie the high flexibility of IPMs and their ability to link different aspects of population ecology, evolutionary biology and life history. The fundamental processes are combined to compute a function called the kernel, which represents all possible transitions between state values through time (e.g. the probability density of transitions from size *x*_1_ at time *t* − 1 to size *x*_2_ at time *t*). The product of the kernel by the number of individuals at time *t* − 1 is integrated over all possible sizes to obtain the number of individuals of size *x*_2_ at time *t*. In general, the numerical implementation of IPMs involves the construction of an iteration matrix to solve the integral. Empirical examples include the study of monocarpic plant species (Childs et al., 2003; Rees et al., 2006; Ellner & Rees, 2006), Soay (*Ovis aries,* Ozgul et al. 2009; Childs et al. 2011) and bighorn (*Ovis canadensis,* Traill et al. 2014) sheep, yellow-bellied marmots (*Marmota flaviventris,* Ozgul et al. 2010), and Trinidadian guppies (*Poecilia reticulata,* Bassar et al. 2016).

A key aspect of the distribution of phenotypes is how traits covary at the level of individuals. Genetic and phenotypic covariances among traits are key determinants of evolution (Lande, 1979). In the context of IPMs, which often consider single traits (e.g. mass), age-specific values of a given trait can be thought of as separate, age-specific traits, the covariances among which are key determinants of evolutionary processes. In fact, in these models, when selection acts only on juveniles, evolution can only occur if there is covariance between trait values at juvenile ages and some aspect (genetic or phenotypic) of the state of individuals at the stage when they reproduce. Such mechanics differ from the classic quantitative genetic approach, where a change in phenotype is transmitted to future generations if there is a change in breeding values. In most IPMs as parameterized to date (e.g. Childs et al., 2003; Ellner & Rees, 2006; Coulson et al., 2010; Ozgul et al., 2010), recovering covariance across ages depends on correctly estimating regressions of observed trait values at each age on trait values at the previous age. In practice, as it is well known, such regressions will typically be underestimated due to regression to the mean (Campbell & Kenny, 1999; Barnett et al., 2005; Kelly & Price, 2005). This statistical phenomenon - unusually extreme measurements being followed by measurements that are closer to the mean - is a manifestation of measurement error. Regression to the mean therefore occurs if phenotypic measurements of predictor variables imperfectly reflect relevant biological quantities. This problem has begun to be investigated in the context of IPMs (Chevin, 2015), and it is likely to be very general. The development functions used so far in IPMs estimate size at any age as an accumulation of growth from birth until that age, which implies that size at age *a* is estimated through a series of regressions in which an increasing measurement error in the predictor is not accounted for. We note that IPMs do not imply the occurrence of regression to the mean. The issues that we discuss in this article are related to the statistical models that are typically - but not necessarily - used in IPMs.

In age-size-structured IPMs, size-dependent transition functions of the fundamental demographic processes are used to project size distribution from one age to the next, and across generations. The inheritance function has been defined as an association between the phenotype of the offspring as newborns or juveniles and that of the parents at the time the offspring was produced (Coulson et al., 2010; Schindler et al., 2013; Traill et al., 2014; Bassar et al., 2016). Essentially, it is a cross-age parent-offspring regression, which is a peculiar measure of resemblance due to inheritance. Outside of the IPM framework, the concept of biometric heritability - the slope of the offspring trait regressed on the midparent’s value (Jacquard, 1983) - is defined by comparing parent and offspring at the same age (e.g. Galton, 1886). In fact, no theory exists for the concept of cross-age heritability as used in IPMs. Body size, commonly the focal trait in IPMs, is typically a dynamic trait (a trait that varies over the development) and therefore its value at a certain age is the result of the accumulation of growth until that age, causing differences among individuals to accumulate over the ontogeny due to environmental and genetic variation in size trajectories (Chevin, 2015). As genes, not phenotypes resulting from development, are inherited, parental phenotype as an adult is an imperfect predictor of the parental genetic contribution to the offspring phenotype. As a consequence of phenotype being used as a predictor, regression to the mean occurs and results in the underestimation of resemblance between parents and their offspring, and therefore of the genetic contribution to phenotypic change (Chevin, 2015).

Here we construct simple but realistic theoretical models of development and inheritance of a quantitative phenotypic trait. For both development and inheritance, we also construct corresponding models to the functions normally implemented in IPMs. By comparing these two sets of models, we investigate how the development and inheritance functions adopted to date in IPMs use data on size-at-age of relatives, and how well they recover across-age and across-generation population structure in continuous traits. Aspects of the distribution of traits through time, other than over single iteration steps, in size-dependent development and inheritance functions, are normally not used to parameterize IPMs. Also, IPMs are typically iterated so that once the population structure at time *t* + 1 is generated, the state of the population at time *t* is discarded. Consequently, while IPMs’ most important feature is tracking the distribution of phenotype through time, they do not output aspects of population structure (e.g. correlations in size within individuals across ages) that allow their performance to be checked. This is a critical point because whenever aspects of the distribution of traits across time are of interest for any inference, particularly evolutionary inference, correlations of individual trait values across ages, and of trait values of relatives across generations, must be adequately reflected. Path analysis (McArdle & McDonald, 1984) can be very useful in studying such correlations. In fact, the structure of both the development functions - with their autoregressive structure - and of the biometric inheritance - with associations both among different generations and among different ages - can be conveniently illustrated by a path diagram representing the causal relationships amongst a set of variables. Also, the path (or tracing) rules are easily applied to obtain the correlations among variables that are not directly associated (e.g. mass at age 1 and mass at age 3). As such, we use path analysis to generate analytical expressions that isolate growth and inheritance, providing insight into the degree to which models of these processes typically used to date recover the structure of populations.

We demonstrate that current parameterizations of IPMs generally recover only a small fraction of the true underlying similarity within individuals across ages (section *Development*), and a small fraction of the true underlying similarity between relatives (section *Inheritance*). These shortcomings have severe consequences for evolutionary inference with IPMs. We then provide an empirical example of a quantitative genetic analysis of developmental trajectories in a pedigreed wild population of bighorn sheep using a random regression animal model of body mass. We compare the random regression analysis, which not only should be robust to regression to the mean, but also uses a model of inheritance based on established principles of how biometric relationships among kin arise from genetic variation (Fisher, 1918; Wright, 1921), to the inheritance function based on the cross-age parent-offspring regression and standard regression methods for growth functions normally implemented into IPMs. We show a large difference between the two parameterizations in the ability to capture similarity within individuals across ages, which results in standard regression methods normally used in IPMs not capturing the across-age structure in growth. Similar conclusions are reached across generations, where IPMs miss most similarity among relatives, corresponding to a failure of the typical IPM inheritance function to predict evolution. We conclude by discussing the results from the theoretical and empirical sections and potential solutions that may prove useful in fully realizing the potential of IPMs.

## Development

Regression to the mean is particularly relevant to IPMs due to how size-dependent growth coefficients are typically - although not necessarily - estimated. Transition rates between size classes for surviving individuals are modelled by regressing observed size at age *a* + 1 on observed size at age *a*, observed size being therefore a predictor. Either linear models (e.g. Childs et al., 2003; Coulson et al., 2010), or extensions of such models, including generalized linear or additive (mixed) models and nonlinear models (e.g. Ozgul et al., 2010; Rees et al., 2014; Traill et al., 2014) have been used for this purpose. All these methods assume that predictors are measured without error. When this assumption is violated, downwardly biased estimates are obtained (for a review on problems and proposed models to deal with measurement error see Thompson & Carter, 2007). Measurements of most traits, including size, will virtually always be made with non-trivial error, for two reasons. First, limitations in the measurement process caused by different measuring conditions (e.g. different levels of stomach fill when measuring the mass of a sheep), or limitations of instruments used for measurement, tend to occur. Second, size, like most other variables of ecological interest, is an abstract concept and therefore is not directly measurable. As such, proxy variables that do not perfectly represent size are measured instead, such as mass or some skeletal measure. The complexity of *size* is such that the covariation between any proxy at time *t* and *t* + 1 is also determined by the other components of size, which are highly correlated with each other. Importantly, the mechanics underlying IPMs neither imply measurement error nor regression to the mean. Rather, the application of standard regression methods that do not account for measurement error within an autoregressive structure on size (subsequent sizes being used as predictors) promotes the occurence of regression to the mean due to measurement error.

Since the measurement error that causes regression to the mean is random rather than systematic, this problem can be modelled by thinking of true size, the trait we want to measure, as a latent variable, *z* that cannot be measured (e.g. McArdle, 2009; Little, 2013, p. 43). In such a scenario, instead of the true values *z*, a proxy, the trait we actually measure, *x*, is recorded, which differs from *z* by a measurement error, 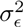, and is related to it by a repeatability, *r*^2^ can, therefore, be written as 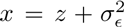. In figure 1A, we illustrate such model of the ontogenetic development of size, which we named *latent true size model*, using a path diagram. In this diagram, true size at age 1, *z*_1_, determines true size at age 2, *z*_2_. *z*_2_ is then a predictor of true size at age 3, *z*_3_, and so on until size at age *n, z_n_*, is predicted. In contrast, the kinds of regression analyses implemented to date in IPMs (e.g. Childs et al., 2003; Coulson et al., 2010; Ozgul et al., 2010; Rees et al., 2014; Traill et al., 2014) assume that true size *z* is being measured when in fact the measured variable is *x*. This model, which we termed *obserevd size model,* is illustrated in Figure 1B. The autoregressive structure in this model is very similar to that in figure 1A, but is built on observed sizes rather than true ones. We use the theoretical models in figure 1 to illustrate the consequences of this conceptual mismatch and to inspect how regression to the mean affects inference about development. We show that the correlations, and therefore the regression coefficients, estimated using IPMs do not correspond to the true latent ones. We then derive a generic analytical expression for how much correlation an IPM can recover given a certain repeatability and number of projection steps (number of IPM iterations).

**Figure 1:**
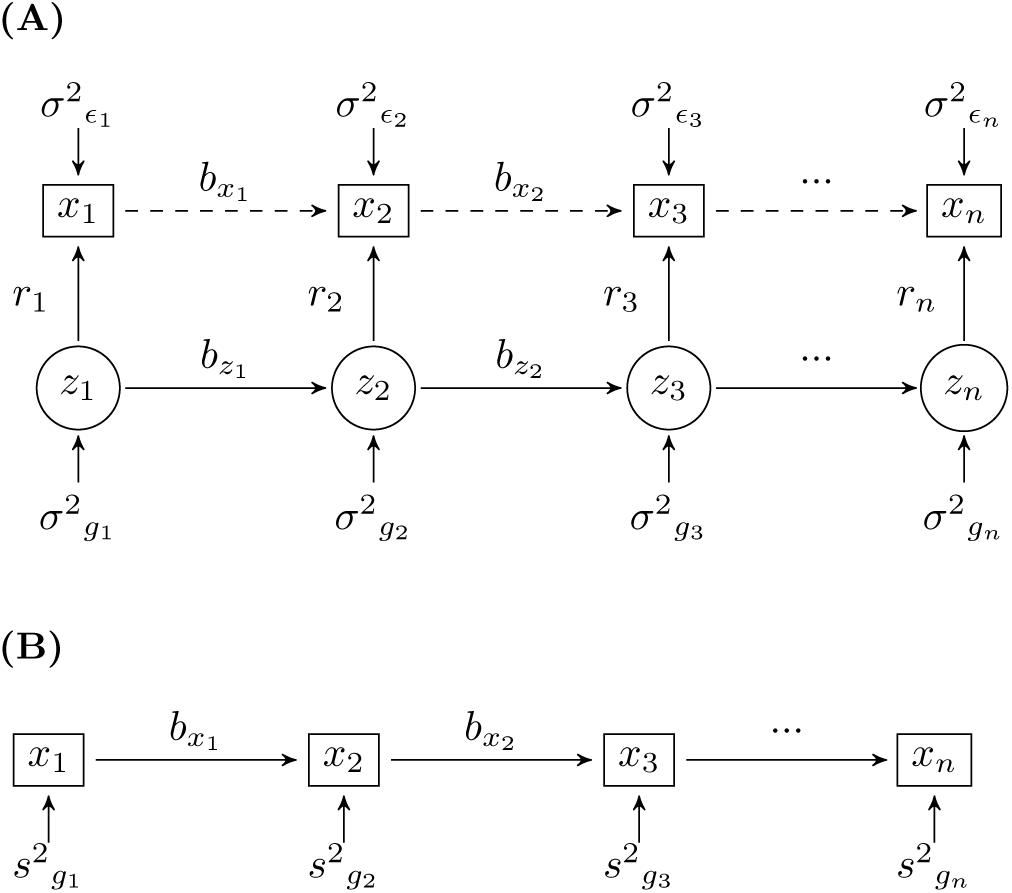
Path diagrams illustrating the ontogenetic development of size. (**A**) Latent true size model; (**B**) Observed size model implemented into IPMs. *z_a_* and *x_a_* are, respectively, the true and observed sizes at age *a*. *r_a_*, linking true and observed sizes, are defined such that repeatabilities are 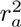. In these antedependence models, 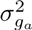 and 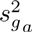 are exogenous variances in growth for true and observed values, respectively, except when they refer to *a* = 1. In this case, 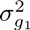 and 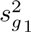 also correspond to variances in size. 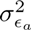 are exogenous errors associated with observed sizes. *b_z_a__* and *b_x_a__* are growth regressions (path coefficients) for true and observed values, respectively. Dashed lines, as opposed to solid lines, do not belong in the path diagram. Although *b_x_a__* correspond to the same quantities in both models, the two models result in covariance structures that are very different (see appendix A.1).

If we consider linear size-dependent growth functions, we can express the true biometric relationships (i.e. true theoretical expressions) among traits *z* (e.g. size at different ages), as well as the relationships captured by standard regression methods typically used in IPMs to describe development, using the principles of path analysis (McArdle & McDonald, 1984). Developed by Wright (1921, 1934) for estimating causal path coefficients, path analysis mathematically decomposes correlations (or covariances) among the variables in a path diagram. For convenience, in the path diagrams that we show we assume that all variables are standardized (mean centered and variance of 1). In such circumstances, the expected correlation between two variables is the product of the standardized path coefficients that link them. Some notational details are worth summarizing: *σ* denotes several aspects of true covariation (covariance in growth among ages), whereas *σ*^2^ represents true variances. Variances estimated by IPMs are denoted by *s*^2^. Since the models in figure 1 are antedependence models (or autoregressive, as the response variable depends on itself at a previous time), 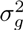 in figure 1A and 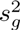 in in figure 1B correspond to variances in growth associated with the regressions of true size on true size at a previous time and observed size on observed size at a previous time, respectively. Finally, the path coefficients *b* correspond to regressions of size on size and *r*^2^ to the square of the regression coefficient of observed size at age *a, x_a_*, on true size at the same age, *z_a_*. Following the principles of path analysis, we used a variance-covariance matrix with the variances in growth, *σ*^2^_*g_a_*_, and errors associated with observed sizes, *σ*^2^_*∊_a_*_ for each age *a*, and a matrix with path coefficients (*b_z_a__* and *r_a_*) matching figure 1A to obtain a variance-covariance matrix for sizes at different ages (Appendix A.1). From this matrix, we then extracted the covariances among ages for both true and observed sizes (Table B.1 in appendix B). As an example, according to the path rules, the correlation and covariance in true size between ages 1 and 3 are given by *b*_*z*1_ · *b*_*z*2_ and *a*^2^_*g*1_ · *b*_*z*1_ · *b*_*z*2_, respectively. Analogous quantities were obtained similarly for IPMs (Table B.2 in appendix B). Since regressions of observed size on observed size, *b*_*x_a_*_, are estimated from the data (rather than implied), these quantities are necessarily recovered correctly, and therefore the *b_x_a__* estimated in IPMs (figure 1B) are equivalent to the analogous quantities in figure 1A. In contrast, variances in growth estimated with observed sizes, *s*^2^_*g_a_*_, do not correspond to variances in growth estimated with true latent sizes, 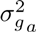, nor to the measurement error associated with observed sizes, 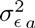. Consequently, since these quantities are crucial to estimate covariances in size among ages, the across-age distribution of phenotype that occurs in a typically-constructed IPM does not generally recover the across-age distribution of either a measured aspect of phenotype (e.g. correlations in the *x* variable across ages) or of an underlying quantity (e.g. correlations in the *z* variable across ages). An across-age distribution of phenotype, which includes correlations among ages, is not typically tracked by an IPM (e.g. Childs et al., 2003; Ellner & Rees, 2006; Coulson et al., 2010; Ozgul et al., 2010). Yet, an IPM’s utility for any ecological and evolutionary inference depends on its ability to track this distribution through time. In a typical implementation, the distribution of phenotype at age *a* − 1 is discarded once the distribution at age *a* is generated, so such correlations cannot easily be outputted and checked against data. As such, we use path analysis to mimic basic IPM mechanics and to extract the across-age dynamics that are not otherwise easily tracked. In contrast, an IPM can easily be interrogated for the distribution of phenotype at any given time. These distributions generally closely match data (Ozgul et al., 2010; Childs et al., 2011, also see fig3(a) in Chevin, 2015 for a simulation example).

For tractability, we demonstrate that IPMs do not in general recover the across-age structure of phenotype using a simplified case of the path diagram in figure 1A as the true model. Specifically, we focus on a static trait, as it renders the basic principles more clearly without loss of generality. We assume that all size-dependent growth coefficients are one (*b_z_a__* = 1, ∀*a*), that the variance in true growth at age one - which also corresponds to the variance in true size at age one - is one (*σ*^2^_*g*1_ = 1) and that the subsequent variances are zero (*σ*^2^_*ga*_ = 0, ∀*a* > 1). Finally, all repeatabilities, *r_a_*, and measurement errors, *σ*^2^*∊_a_*, take the same value, *r* and 1 − *r*^2^, respectively. Applying the path rules and these assumptions results in the particular case of all true phenotypic variances and covariances being 1 and variances and covariances for phenotypic observed size being 1 and *r*^2^, respectively (see appendix A.1.2 for details). Standard regression methods typically used in IPMs underestimate regressions for true growth in any instance where *r* < 1, by a factor of *r*^2^. Whenever true and observed sizes differ, which is true for virtually every attempt to measure size, instead of 1 (value set for all *b_z_a__*), *b_x_a__* take the value *r*^2^ for any consecutive pair of ages (both in figure 1A and 1B). As mentioned before, covariances in size across ages are in general not reported when building an IPM. However, the implied covariances can be calculated using path analysis (see appendix A.1.1 for the general case and appendix A.1.2 for this simplified example). Since according to the path rules of standardized variables correlations between two variables correspond to the product of the path coefficients linking them, in this example correlations in size among two ages will be *r*^2^ to a power equivalent to the number of links between them. As such, since *r* < 1, these correlations will be underestimated. As for the covariances, these are obtained by multiplying the correlations by the variance in growth at age 1, which corresponds to the variance in growth at age one, *s*_*g*1_. Variances in size are well recovered in IPMs because these quantities are directly estimated from the data. Therefore, in this example, *s*_*g*1_, which also corresponds to variance in size at age one, corresponds to one, resulting in covariances in size implied by the growth functions normally implemented in IPMs being given by

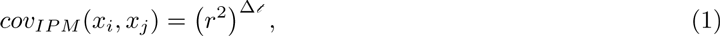

where Δ*𝓉* is the number of projection steps (or path coefficients) connecting ages *i* and *j* (*j* − *i*).

The standardized conditions set in this simplified example illuminate how much correlation between sizes at different ages the standard IPM formulation will miss. As true correlations (or covariances) in size across ages were set to one, subtracting the correlation in equation (1) to that theoretical value corresponds to the amount of correlation a standard regression fails to recover,

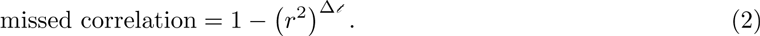

The theoretical result of equation (2) shown in figure 2 demonstrates that this quantity is far from negligible, increasing rapidly with the number of projection steps and decreasing values of *r*. Many IPM analyses to date have focused on long-lived organisms. In these systems, age differences (projection steps) of 5 to 10 years may correspond to the gap between juvenile stages, which are often subject to the strongest viability selection, and ages of greatest fecundity. Even for traits with high repeatabilities (e.g. *r* = 0.9), correlations over such age differences will be underestimated by more than 60% (Figure 2). Ultimately, size is estimated as an accumulation of growth through an autoregressive process that discards the distribution of size at time *t* − 1 at each iteration (when the distribution at *t* is obtained). This results in measurement error at each iteration not being accounted for in the next, and therefore the effect of regression to the mean rising with the number of IPM projection steps. Serious consequences can be expected both for evolutionary and ecology studies, whenever differences in individual growth are of interest. Curiously, all else being equal, IPMs with narrower projection intervals (e.g. monthly, rather than yearly) will suffer more from regression to the mean than models constructed with wider projection intervals. Finally, it is important to note that asserting that the observed quantities, rather than underlying variables, are the target of interest in any given IPM application does not solve the fundamental problem. In any scenario where the covariance of observed values through time is caused (in part or in whole) by quantities other than the observed values themselves (figure 1A) a model of sequential regressions of observed values on one another (figure 1B) will not recover the resulting covariance structure.

**Figure 2:**
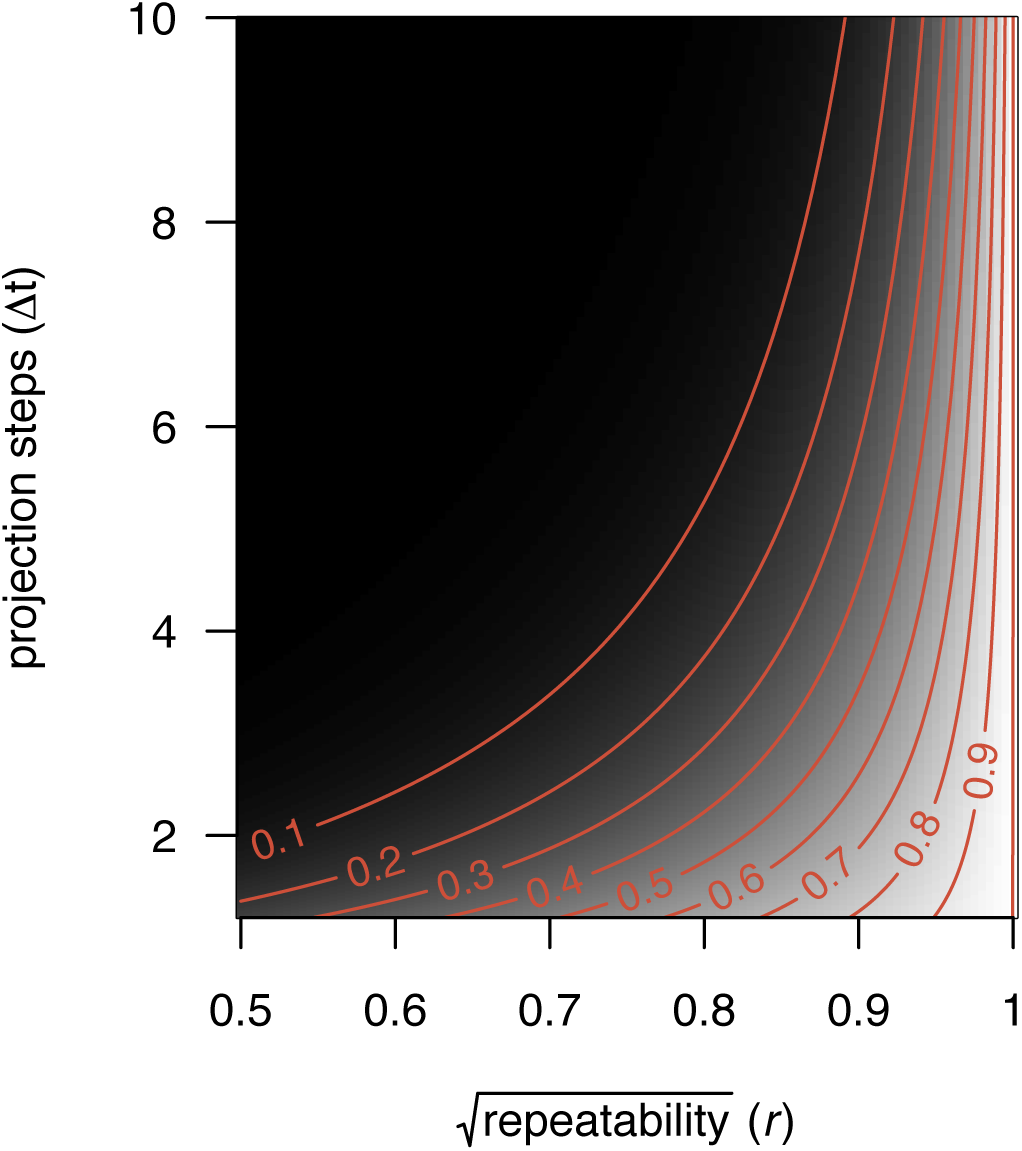
Proportion of correlation in size among ages recovered by a typically-built IPM as a function of the square root of the repeatability (*r*) and number projection steps (Δ*𝓉*). The true values were used as reference.

## Inheritance

The modern understanding of how genes contribute to similarity among relatives (Fisher, 1918, 1930; Wright, 1922, 1931) has a very different structure from the inheritance function typically included in IPMs (e.g. Coulson et al., 2010; Traill et al., 2014; Bassar et al., 2016). Fisher and Wright showed how Mendelian inheritance at many loci influencing a trait generates the observed biometric relationships among relatives, including the relationships of a quantitative character between parents and offspring. Here, we use the basic mechanics of inheritance of a polygenic trait, which have well-known relationships to selection and evolution (Walsh & Lynch, forthcoming), and use it as simple background to see if IPM mechanics are generalizations of these principles. The notion of breeding value, or genetic merit, of an individual is central to the current theory of the inheritance of quantitative traits, and has its roots in Fisher’s (1918) and Bulmer’s (1980) infinitesimal model (see Falconer, 1981; Walsh & Lynch, forthcoming, Chapter 15). Each parent passes half of its genes and therefore half of its breeding value on to the offspring. As such, the expected breeding value of offspring *i*, 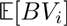, corresponds to half the sum of parental breeding values, as follows

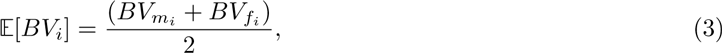

where *BV_m_i__*. and *BV_f_i__*. are the maternal and paternal breeding values, respectively. The true breeding value, *BV_i_*, follows a normal distribution,

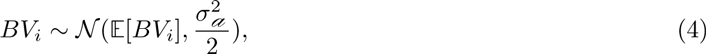

with its expected value as mean and 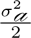 as variance, corresponding to the variance in breeding values in the absence of inbreeding, conditional on mid-parent breeding values, resulting from segregation (Bulmer, 1980). The variance in the breeding values divided by the phenotypic variance is defined as heritability, *h*^2^, a measure of evolutionary potential. The degree of resemblance between relatives provides the means for distinguishing the different sources of phenotypic variation and therefore for estimating heritabilities and other quantitative genetic parameters (Falconer, 1981). The simplest way of doing so is by using correlations of close kin, for example, of parents and their offspring, as *h*^2^ corresponds to the slope of the offspring trait regressed on the midparent’s (Lynch & Walsh, 1998, Chapter 7). In fact, Jacquard (1983) defines the heritability estimated with a parent-offspring regression as a biometric heritability, as opposed to broad- and narrow-sense heritabilities, for which the genetic and additive genetic variances are, respectively, explicitly estimated. Any genetic architecture, i.e. broad- and narrow-sense heritability, determines the biometric relationships among kin (Lynch & Walsh, 1998, Table 7.2). In IPMs, heritabilities have been estimated using parent-offspring regressions. Specifically, inheritance has been defined as a regression of the phenotype of the offspring as newborns or juveniles on that of the parents at the time the offspring was produced (Coulson et al., 2010; Schindler et al., 2013; Traill et al., 2014; Bassar et al., 2016). In this section, we investigate whether this cross-age biometric notion of inheritance is compatible with what is known about trait transmission across generations.

### Inheritance across generations

We start by addressing the consequences of regression to the mean related to the biometric concept of inheritance when it is applied across multiple generations. We defined a true model for trait transmission across four generations of the same age, according to Fisher’s and Wright’s understanding of trait transmission (Figure 3A), and a comparable model reflecting the biometric concept of inheritance typically used in IPMs (Figure 3B). As for the development models, we used path diagrams and path analysis to compare the correlations implied by both models. In figure 3A, breeding values, the underlying units that are inherited, are passed on across generations: from great-grandparents to grandparents, from grandparents to parents, and from these to the offpring). Since each parent passes on half its breeding value to the next generation, the regression coefficient linking generations is 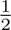. The variance associated with the breeding values is 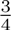, which corresponds to 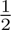 from the other parent and 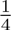 from segregation. *h* corresponds to correlation between the breeding values and phenotypic values (Wright, 1921; Falconer, 1981) and, in a standardized path analysis, to the corresponding regression coefficient as well. If observed size is standardized (variance of 1), then according to the path rules its exogenous variance corresponds to 1 − *h*^2^. Finally, if any regression was to be made between the observed sizes, *x*, the coefficient would be half the heritability. There is a close analogy with the path diagrams in figure 3A and figure 1A. Not only do they share the same structure (sizes at different generations instead of sizes at different ages), but other analogies can be taken. For example, as the regression coefficient of phenotype on breeding values, the square root of the heritability expresses the reliability of the phenotype to represent the underlying genetics, which in figure 1A was represented by the square root of the repeatability. In figure 3B we show a series of parent-offspring regressions based on phenotype, rather than genetics. The slope of the parent-offpring regression for a single parent is known to be 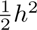 and in a standardized path analysis, the associated variance is 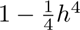. Similarly, the path diagram in figure 3B relates to the one in figure 1B.

**Figure 3:**
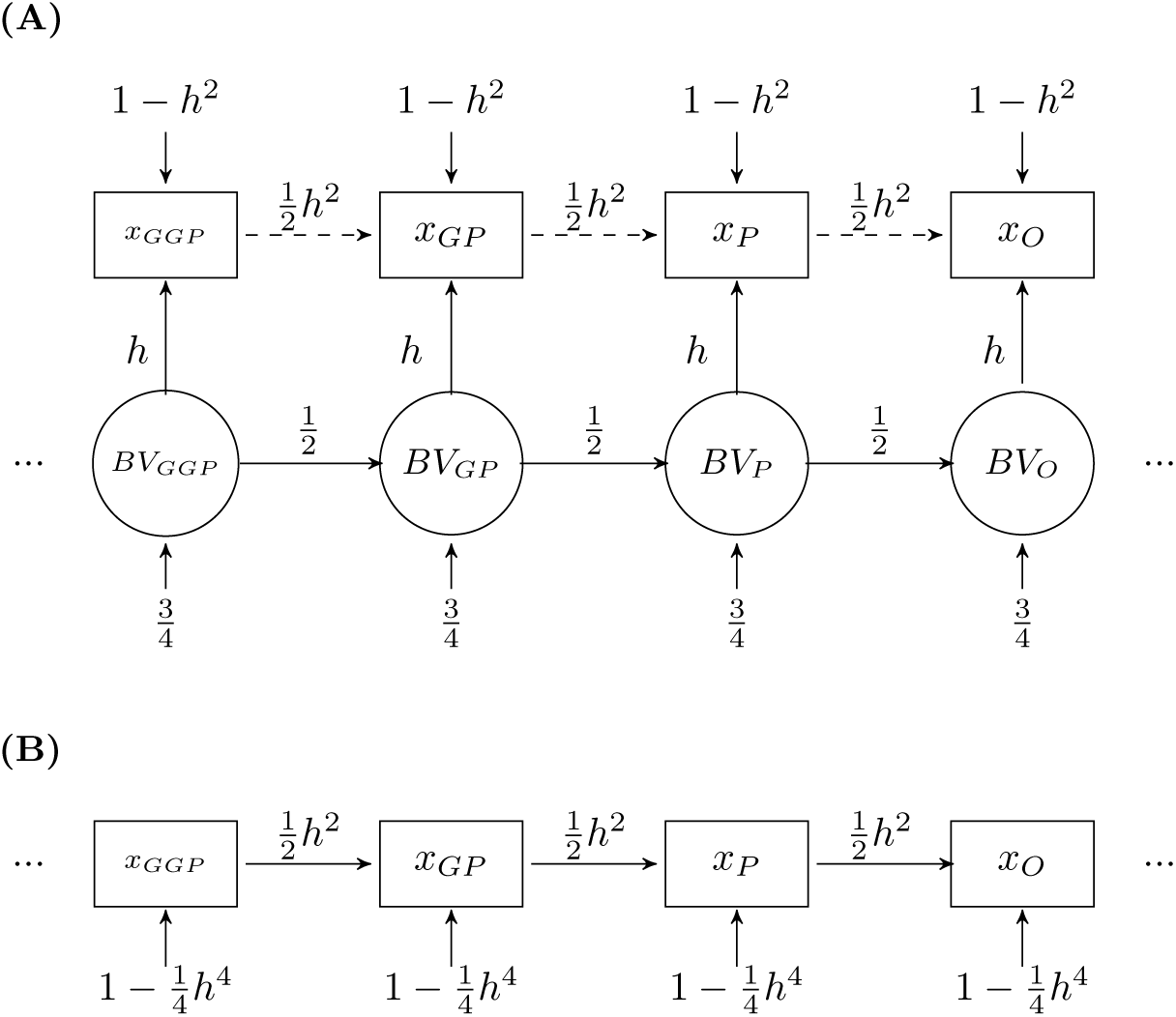
Path diagrams illustrating the transmission of a quantitative trait across generations of the same age. (**A**) Model based on the fundamentals of quantitative genetics; (**B**) model corresponding to a purely biometric notion of inheritance. *BV* and *x* correspond to breeding values and the observed phenotype, respectively. The exogenous inputs to BVs include contributions from the other parent and segregation. The subscripts *GGP*, *GP*, *P* and *O* denote great-grandparent, grandparent, parent and offspring, respectively. *h*^2^ corresponds to the heritability and therefore *h* and *h*^4^ to its square root and square, respectively. Dashed lines, as opposed to solid lines, do not belong in the path diagram. While the observed parts of the two models look very similar, they imply different correlation structures among relatives more than one generation apart (see main text).

With this single age set up, we can isolate the regression to the mean that occurs as a result of a purely biometric approach to the inheritance function. As for the true regressions, parent-, grandparent-, great- grandparent-offspring regressions are given by 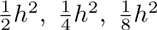, respectively (Lynch & Walsh, 1998). The extension for arbitrary ancestral regressions is given by

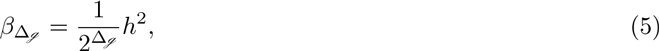

where Δ_*ℊ*_ is number of generations between two relatives. We used path analysis to obtain the analogous regressions that are implied when applying a biometric inheritance function repeatedly within an autoregressive process (Figure 3B). The structure of the path diagrams in figures 1B and 3B are equivalent and therefore the reasoning for obtaining covariances and regressions for size presented in appendix A.1 also applies in this case. As such, according to the path rules, IPMs, as usually parameterized, will estimate these regressions as

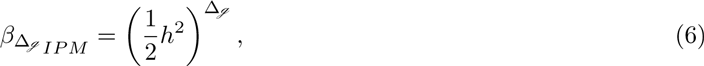

which does not correspond to equation (5). As an example, tracing the regression of grandoffspring size (*x_O_*) on grandparent size (*x_GP_*) in this standardized path diagram involves two paths with coefficient 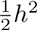, resulting in 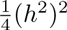 instead of the true regression 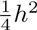. Equation (6) implies that trait transmission between same-age relatives is not fully recovered when the gap between generations (Δ_*g*_) is greater than one. For ancestral regressions other than of offspring on parent to be correctly recovered the heritability of this trait would have to be one, which tends not to happen in nature for most ecologically interesting traits. The proportion of the true regressions recovered by the biometric inheritance function is given by *h*^2(Δ_g_−1)^, as illustrated in figure 4. For example, if a trait has a heritability of 25%, the grandparent-grandoffspring regression will be estimated as 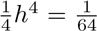 rather than its true value of 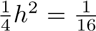, which corresponds to only recovering 25% of the regression. This proportion drops to 6.25% for great-grandparents and their offspring.

**Figure 4:**
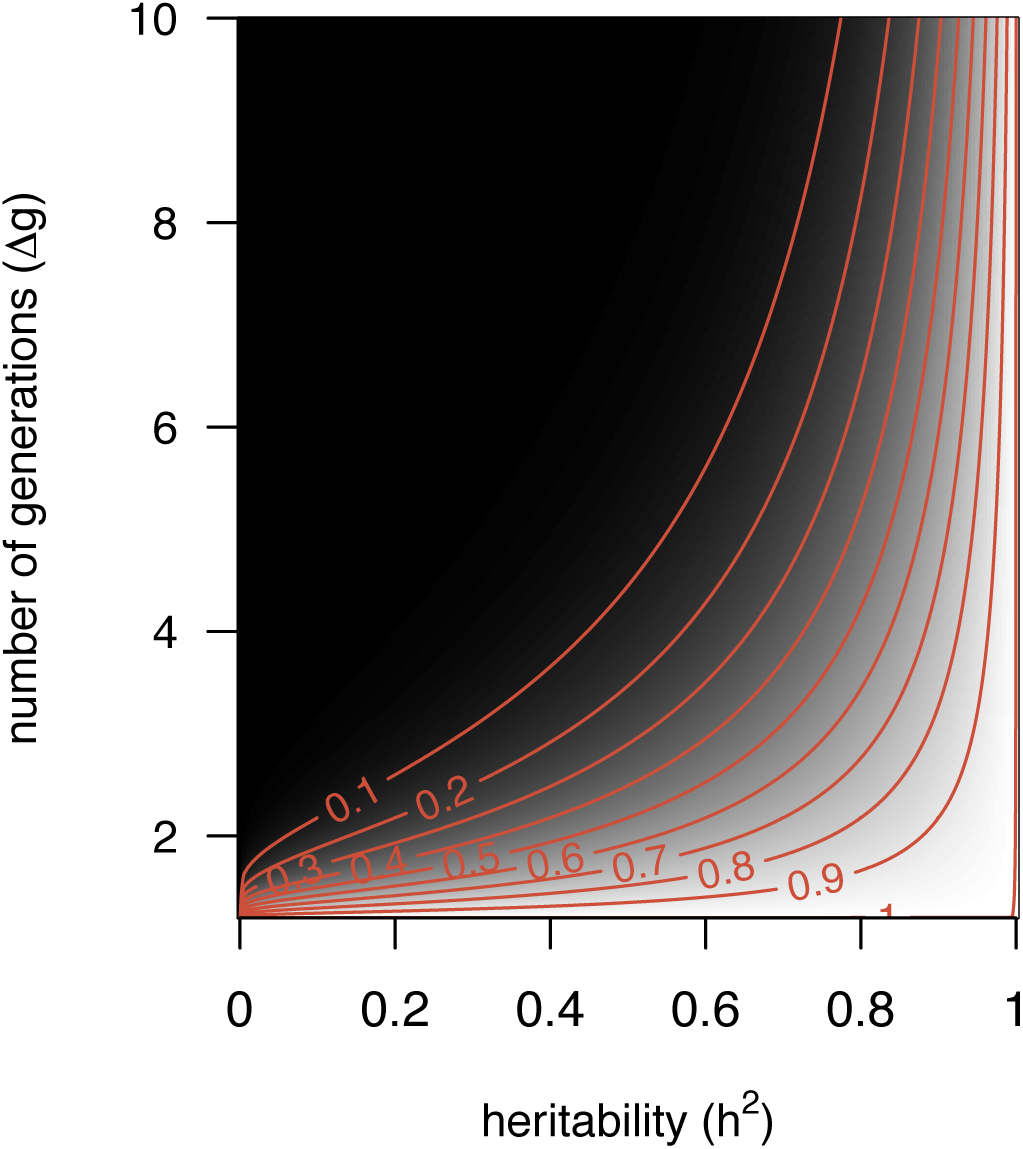
Proportion of the parent-offspring regression recovered by a same-age inheritance function as a function of the heritability (*h*^2^) and the number of generations (Δ_*ℊ*_). The true values were used as reference.

### Across-age inheritance functions

There is a second mechanism by which regression to the mean affects inference with the inheritance function, particularly resulting from its cross-age structure. It is important to note that although an individual’s genetic constitution is constant throughout life, the genetic variants relevant at one life stage need not affect other life stages. Genetic variants acting late in life may be latent early in development. Such variants may be inherited and contribute to similarity among relatives, even if they contribute neither to covariance of traits within individuals, through time, nor to covariance of parents, as adults, with their offspring, at young, or arbitrary, life stages. Consequently, there is potential for the concept of inheritance applied to date in IPMs to neglect a major fraction of how genetic variation can generate similarity among relatives (Hedrick et al., 2014; Chevin, 2015). Chevin (2015) illustrated this issue with numerical demonstrations. Here we formalize his findings analytically to explore the generality and the magnitude of his conclusions. We examine what would happen to two cohorts (parents and offspring) with two ontogenetic stages (juvenile, J, and adult, A, Figure 5). We choose a simple model with only two ontogenetic stages, since extending it to include more age classes would correspond exactly to what was described for development in the previous section. We explore two different perspectives of trait transmission - first using basic quantitative genetic principles and then a cross-age biometric approach typical of IPMs. The first path diagram (Figure 5A) reflects the former, with phenotype being a result of the breeding values, BV, and the environment, *σ*_ℯ_^2^. To account for the fact that different genes may influence different traits or the same traits across ages, we use different symbols for breeding values in the juvenile and adult stages. In this path diagram, parent phenotype as a juvenile determines parent phenotype as an adult through the regression coefficient *b*. We also represent segregation and mating, through which the offspring receives paternal breeding values that, together with the environment, define offspring phenotype as juveniles, *O_J_*. Finally, offspring phenotype as juvenile also determines its phenotype as an adult, *O_A_*. We use the subscripts *z*, *a* and *e* to distinguish between phenotypic variance, *σ*^2^, and covariance, *σ*, and their additive genetic and environmental components, respectively. The diagram in figure 5B illustrates a cross-age phenotypic transmission between parents and offspring normally used in IPMs (e.g. Coulson et al., 2010; Traill et al., 2014; Bassar et al., 2016). In this diagram, parent phenotype as a juvenile determines parent phenotype as an adult (through the regression coefficient for development, *b_dev_*), which determines offspring phenotype as a juvenile (through the regression coefficient for inheritance, *b_inh_*). Finally, growth also occurs in the offspring, resulting in its adult stage. As before, we consider linear size-dependent growth functions, and additive genetic effects on juvenile size and subsequent growth, so that path analysis can be used to obtain the biometric relationships among traits (true theoretical expressions), as well as the relationships captured by the cross-age inheritance function implemented in IPMs (see appendix A.2 for details). First, we defined true hypothetical additive genetic and environmental variance-covariance matrices for growth at each age, as well as true path coefficients that match the path diagram in figure 5A. Subsequently, we used path analysis to obtain the true phenotypic variance-covariance matrix for size, a matrix that quantifies both direct and indirect effects of size at each age. Finally, the slopes of the regressions of offspring size on parent size were obtained analytically from the model, corresponding to the true parent-offspring regressions for both juveniles,

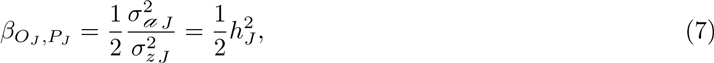

and adults,

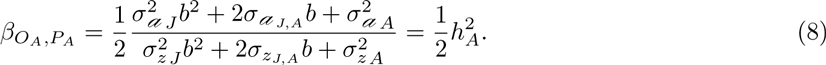

**Figure 5:**
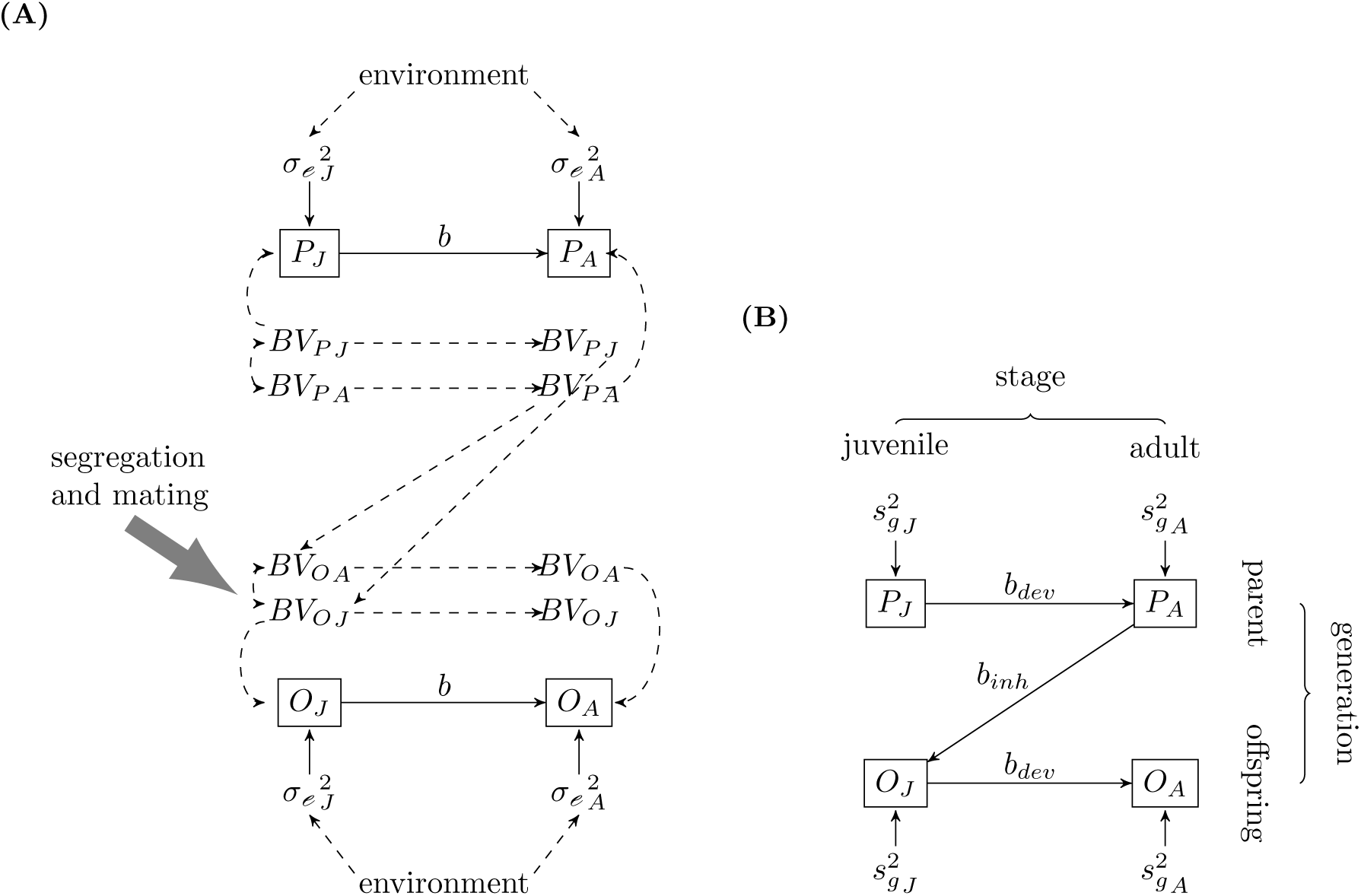
Path diagrams illustrating the transmission of a quantitative trait between parents, *P*, and offspring, *O*, with two ontogenic stages, juvenile, *J*, and adult, *A*. (**A**) Model based on the fundamentals of quantitative genetics; (**B**) model corresponding to a cross-age concept of trait transmission. *P_J_* and *P_A_* correspond to parental trait as juvenile and adult, respectively, and likewise for the offspring (*O_J_* and *O_A_*). 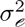 and 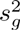 correspond to the exogenous variances of size at birth, and of growth until the juvenile stage (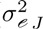 and 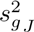) and of growth (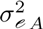 and 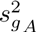). *b, b_dev_* and *b_inh_* correspond to regressions, namely for development (*b* and *b_dev_*) and inheritance (*b_inh_*). Finally, *BV* are breeding values. Although the genetic constitution is constant over an individual’s life, different genes are activated throughout life, which is denoted by distinguishing *BV* for both juvenile and adult stages.

Note that the numerator and denominator in equation (8) are simply reconstructions of the additive genetic and phenotypic variances in size, respectively, given the additive genetic and phenotypic variances in juvenile size, growth to adult size, and the covariance between them. Two other expressions are required, as they are used in constructing IPMs, namely for the regression of adult offspring size on juvenile offspring size, or adult parent size on juvenile parent size,

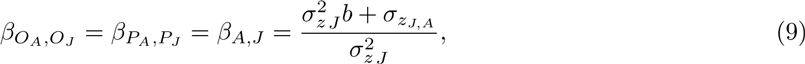

which models the ontogenetic development of size, and for the regression of juvenile offspring size on adult parent size,

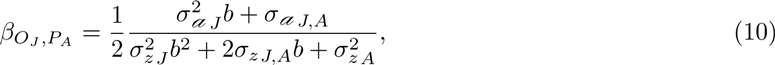

which corresponds to the cross-age inheritance function.

As shown in figure 3B, typical IPMs adopt *β_O_J_, P_A__* (*b_inh_*) as the inheritance function. We use the path rules to obtain the covariances among same-age parent and offspring that are implied by this quantity, and therefore to obtain expressions for the same-age parent-offspring slopes. In practice, we then compare the theoretical results presented above, in particular the true parent-offspring regressions in equations (7) and (8), to those that occur with the cross-age inheritance function, allowing us to derive the conditions under which IPMs recover the population structure of continuous traits between parents and offspring. According to the path rules, IPM-based inference for parent-offspring regression at both juvenile and adult stages, *β_O_J_, P_J__* and *β_O_A_, P_A__*, respectively, corresponds to the product of *β_J, A_* (Equation 9) and *β_O_A_, P_A__* (Equation 10, see appendix A.2 for details), as follows

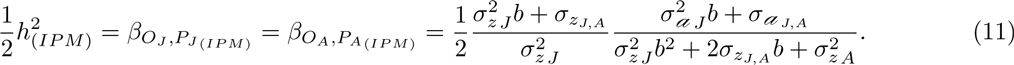

As a result, in a two-stage case, an IPM as typically built implies the same value of the parent-offspring regression for both stages, which is not the case for the true values (Equations 7 and 8). Also, and even more importantly, the IPM-based inference corresponding to the expression in equation (11) does not correspond to the true values for either age (Equations 7 and 8). Thus, IPMs do not, in general, recover parent-offspring regressions.

The comparison between IPM-based inference and true values becomes more straightforward in the simplified case of no covariances of growth across ontogenetic stages (additive genetic, *σ_𝒶_J, A__*, and more generally, phenotypic, *σ_z_j, A__*), which is assumed in all IPM implementations to date for developing traits. In such circumstances, the IPM implies a parentoffspring regression, for both juveniles and adults, of

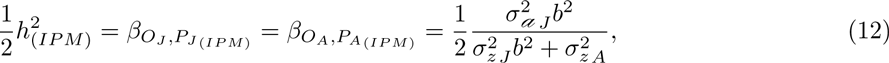

which is always less than the corresponding true values. This is a best-case scenario for IPMs, as covariances of growth across ages are in general not modelled when estimating size transitions in such models. Even in such unrealistic conditions, a standard IPM can only recover the true parent-offspring regressions under very specific conditions. According to equation (12), for parent-offspring regression in juveniles to be fully recovered by a model using a cross-age biometric inheritance function, the phenotypic variance in growth, 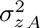, must be zero. When that is not the case, the proportion of regression recovered decreases with decreasing size-dependent size regression, *b* (Equation 7, Figure 6A). The same condition holds for the parent-offspring regression in adults (Equation 8, Figure 6B). These quite narrow conditions are unlikely to occur in nature. We obtained similar results for the case where covariance in growth exists (Appendix B). Indeed, although IPMs were developed to model dynamic traits, the conditions for which they are guaranteed to recover parent-offspring regression, particularly the absence of variance in growth, essentially constrain a dynamic trait to be static.

**Figure 6:**
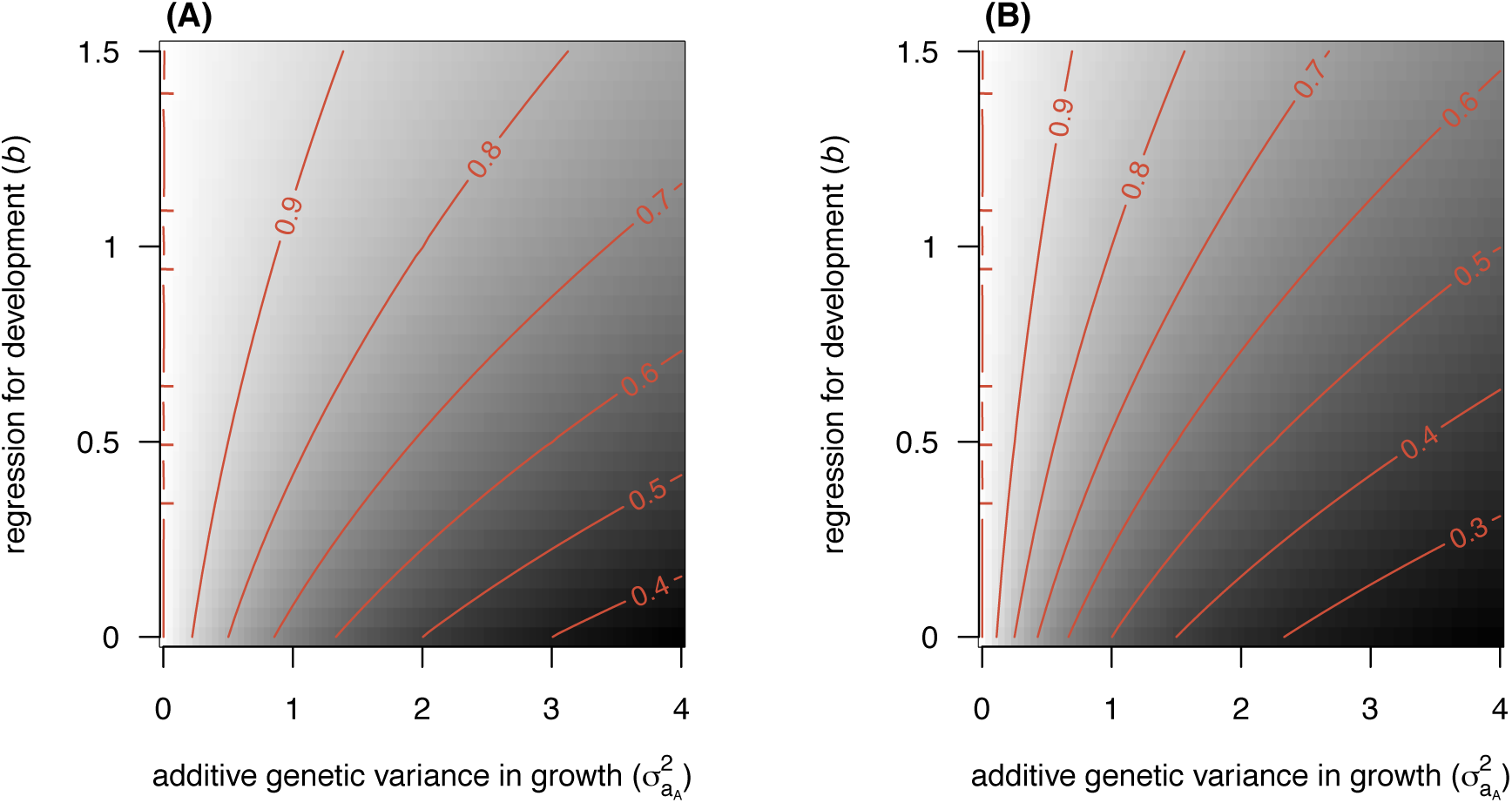
Proportion of parent-offspring regression recovered by a cross-age parent-offspring regression, in juveniles, (**A**), and adults, (**B**). In both cases, correlation in growth, genetic (*σ_a_J, A__*) and environmental (*σ_𝒶_J, A__*), was assumed to not exist, and the remaining parameters were set as follows 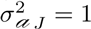, 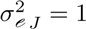, and 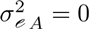. The true values were used as reference in both plots.

### Parent-offspring regression with a constant intercept

The preceding analysis shows that regression to the mean prevents the inheritance function from capturing most aspects of covariance between individuals and their descendants. In language typically used to describe properties of IPMs, a cross-age biometric inheritance function does not fully capture the most important ways in which inheritance influences the dynamics of a population through time. Importantly, however, as shown above, the biometric inheritance notion does capture the correct covariance of parents and offspring, at least of a static trait (or a model with a single age class). In itself, this may imply that a purely biometric notion of inheritance can be used, at least in simple cases, to track some important features of a population. Nonetheless, the use of the concept of biometric inheritance that is extensively recommended for IPMs (Coulson et al., 2010; Coulson, 2012; Rees et al., 2014) does not correctly employ the concept. This recommendation is based on two misconceptions about biometric inheritance, both of which lead to failures to characterize even the simplest aspects of phenotype (e.g. the dynamic of mean phenotype). The first misconception, shown above, is the assumption that theory underlying the biometric relations among kin can be applied to a non-static trait when parents and offspring are of different ages. This includes the assumption that iteration of the purely phenotypic relations of parents and offspring across multiple generations can recover biometric relationships among more distant kin, e.g. arbitrary ancestral regressions. The second misconception is that the biometric inheritance concept, and its known relationships to quantitative genetic parameters (Lynch & Walsh, 1998, Chapter 7), implies that biometric functions are constant. A constant genetic basis (e.g. an assumption that *h*^2^ is constant over a period of time) to a trait is commonly assumed in quantitative genetic studies, and implies that the slope of the parent-offspring regression is constant. However, should a trait evolve, changing the mean phenotype, then the intercept of the parent-offspring regression necessarily changes. If the intercept is assumed to not change, or a model is constructed where the intercept cannot change, then the dynamic of mean phenotype will be highly restricted. Therefore, even the simplest possible IPM constructed with a typical inheritance function, which has not only a constant slope, but also a constant intercept, will necessarily fail in describing the evolution of mean phenotype.

As an example, consider a non-age structured population, with no class structure other than that associated with some focal trait, *z*. We denote the mean trait value in generation ℊ by *Z̄*ℊ and its heritability as *h*^2^. Without loss of generality, we assume that during a period of equilibrium *z* is measured such that its mean is 10. We also assumed that *z* is heritable (*h*^2^ = 0.5) but, since there is no selection, no phenotypic change is observed (Figure 7A). Suppose that the equilibrium is then disrupted and that both sexes experience the same selection, which represents a change in mean phenotype for the first generation 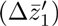 of 1 unit (Figure 7B). The offspring on mid-parent regression is then 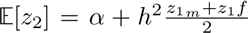, where a is the intercept and *z*_1_*m*__ and *z*_1_*f*__ denote maternal and paternal phenotypes, respectively. An IPM constructed using this regression (appropriately handling the two sexes) yields a mean phenotype in the next generation of 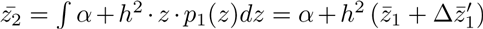. The first expression corresponds to the integral that would be solved (typically numerically) by an IPM corresponding to this example, and *p*_1_(*z*) is the probability density function of phenotype after selection but before reproduction in generation 1. The second expression is the analytical solution for this integral, made possible by assuming a linear function. Under the conditions set for this example, this expression would be 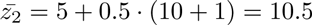. This change satisfies the breeder’s equation for the change in mean phenotype across generations 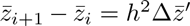. The problem arises in the next generation.

**Figure 7:**
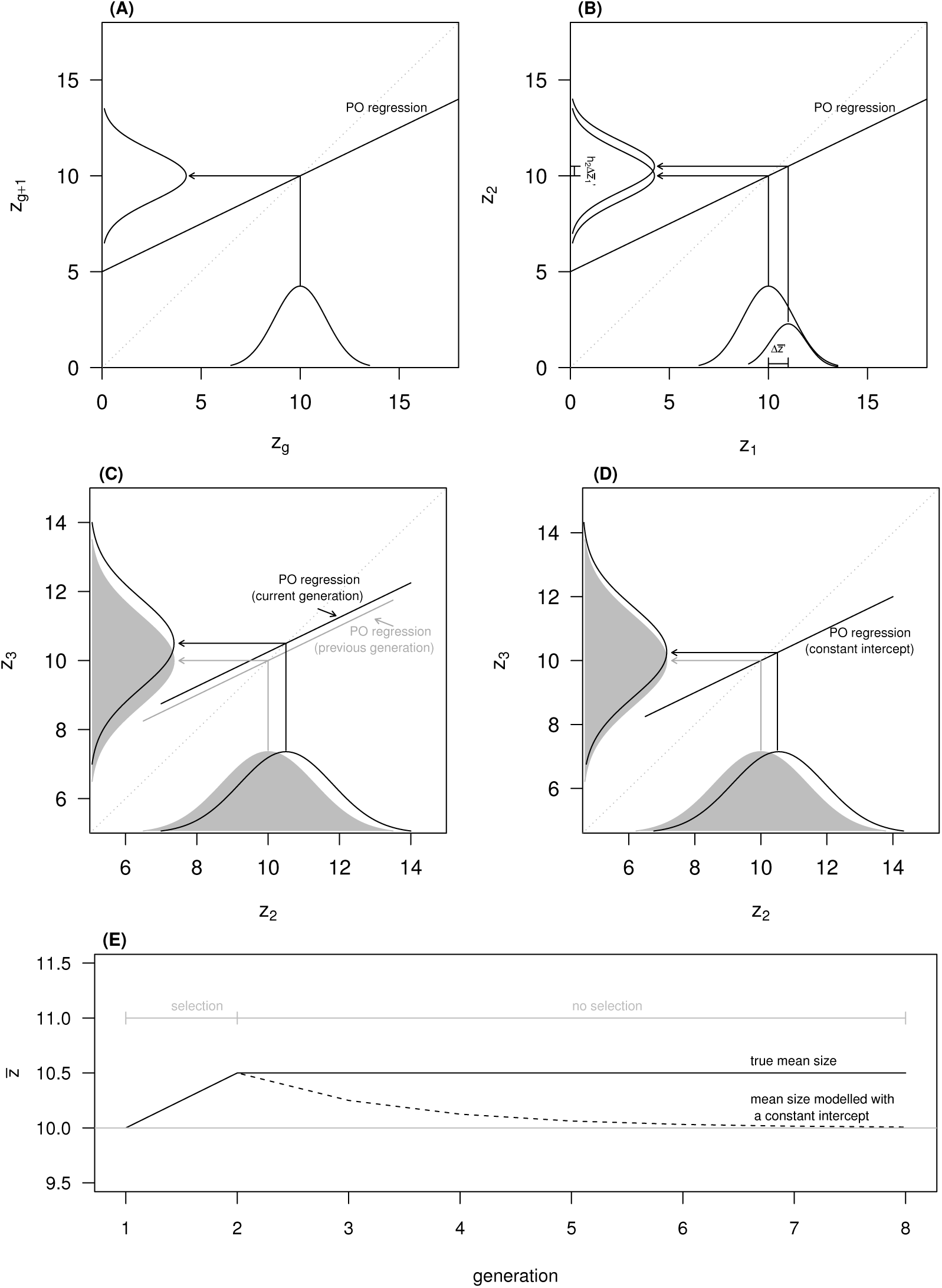
The consequences of assuming a constant intercept for the parent-offspring (PO) regression across generations. (**A**) Population at equilibrium, where mean phenotype is 10; (**B**) Period of selection. Selection before reproduction causes mean parental size to change from 10 to 11 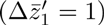. Mean offspring phenotype (*z*_2_) is 10.5, which implies a parent-offspring regression given by 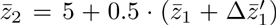, and therefore *h*^2^ = 0.5 and an intercept of 5; (**C**) Relaxed selection. When mean phenotype changes across generations, in this case from 10 to 10.5, the intercept of the parent-offspring regression necessarily changes as well. In a case of no selection in generation 2, the parent-offspring regression is given by 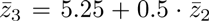; (**D**) Relaxed selection with constant intercept. If the intercept is assumed to remain constant, and the first parent-offspring regression is used to estimate the mean phenotype in generation 3 (*z*_3_), instead of the true value 10.5, 10.25 is obtained instead; (**E**) Iteration of mean phenotype to subsequent generations of relaxed selection, both under a model with a genetical notion of inheritance and an analytical iteration of a simple IPM with a biometric inheritance function with a fixed intercept. In (**C**) and (**D**) the distribution in grey corresponds to the previous generation.

Let us suppose that selection is now relaxed, such that the within-generation change in phenotype due to selection, 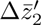 is zero. In the absence of selection, drift, immigration and mutation, we expect no change in allele frequencies (Wright, 1937) and therefore no evolution. Consequently, we expect no change in mean phenotype (Figure 7C). In a very simple non-age structured IPM, we would use the current distribution of trait values (*g* = 2) and the same inheritance function to obtain the mean phenotype in generation 3, and that would correspond to 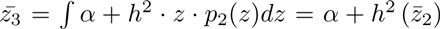, which in this case would be 10.25 (Figure 7D). In this example, an IPM would predict the trait moving back 0.25 phenotype units, which corresponds to reverting back to half of the initial response to selection. If *z*_2_ is any value other than 10, the static biometric inheritance function results in changes in mean phenotype in the absence of selection, drift, mutation and migration. Continuing the analytical iteration of the mean phenotype in this simple IPM, we show that with each subsequent generation (iteration step, in this simple argument), the mean phenotype regresses further toward a value determined by the nature of the static biometric inheritance function (Figure 7E). If selection is sustained, then the dynamic of the mean phenotype even in this very simple IPM will be wrong, representing a component associated with the response to selection, and a spurious change due to the misconception of biometric inheritance associated with a parent-offspring regression with a fixed intercept. A biometric inheritance function with a constant slope and intercept is inconsistent with evolution.

### Study case: bighorn sheep

We used a pedigreed population of bighorn sheep from Ram Mountain, Alberta, Canada (52°N, 115°W) to assess the performance of the development and inheritance functions as implemented in standard IPMs. Both quantitative genetic (e.g. Coltman et al., 2003; Wilson et al., 2005) and IPM analyses (Traill et al., 2014) have been conducted for this study system. This isolated population has been the subject of intensive individual-level monitoring since 1971. Sheep are captured and weighed multiple times per year between late May and late September. For detailed information on the study system see Jorgenson et al. (1993), Festa-Bianchet et al. (1996) and Coltman et al. (2003). We analyzed individual age-specific masses adjusted to September 15 (see Martin & Pelletier, 2011; Traill et al., 2014) for 461 ewes captured from 1975 to 2011 and aged up to 10 years (2002 ewe-years). We built two statistical models, one reflecting how the ontogenetic development of size and inheritance have been typically modelled in IPMs, and the other corresponding to a possible alternative to estimating these two key functions, a random regression animal model of body size (Kirkpatrick et al., 1990, 1994; Meyer & Hill, 1997; Meyer, 1998; Wilson et al., 2005). We chose random regression because it is widely used to study the genetics of developmental trajectories and it satisfies a number of criteria, namely: (i) it accommodates across-age covariance, over and above that attributable to measured values of focal traits, (ii) it incorporates the known fundamentals of quantitative genetics, (iii) it is economical in terms of the number of parameters that need to be estimated, and (iv) its basic structure is compatible with IPMs. Criteria (i) and (ii) result in random regression analysis providing an approach for characterizing development and inheritance that should be robust to regression to the mean, as imperfectly measured quantities are not used as predictor variables, and as it uses a modern notion of inheritance of quantitative traits. Nonetheless, other options can also avoid regression to the mean, including a formulation of an explicit genetic autoregressive size-dependent model that accounts for measurement error. Also, although the random regression approach, and potentially other models using quantitative genetic approaches characterizing variation in phenotype and its inheritance, could profitably be integrated into the broader IPM framework, for simplicity we refer to the former approach as “IPM” and to the latter as “RRM”. Both models were fitted in a Bayesian framework, using MCMCglmm (Hadfield, 2010), and diffuse inverse gamma priors for all (co)variance components.

### Standard IPM approach

We used a linear model to estimate the development and inheritance functions used in typical IPMs. We modelled observed ewe mass at each age as a function of mass at the previous age, with separate intercepts and slopes for each age. For lambs, we estimated a regression of lamb mass on the mass of their mother two months before conception (previous September). Formally, the model is described as

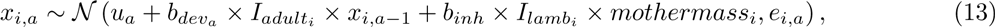

where *x_i, a_* is the observed mass of individual *i* at age *a*, *u_a_* age-specific intercepts, *b_deu_a__* age-specific size slopes and *b__inh__* is the inheritance function coefficient. *I_lamb_* and *I_adult_* are indicator variables for lambs and older individuals, respectively. Finally, *e_a_* are heterogeneous residuals per age. The estimated fixed effects and variance parameters are presented in table 1.

**Table 1:** Coefficients for the IPM standard approach, including regressions of mass at age *a* on mass at age *a* − 1, and of lamb’s mass on mother’s mass at conception for the bighorn sheep population of Ram Mountain. The values correspond to posterior modes and 95% quantile-based credible intervals.

| Age | Intercept | Slope | Residuals |
| --- | --- | --- | --- |
| 1 | 17.37 ( 14.14 - 20.59 ) | - ( - - - ) | 17.80 ( 15.97 - 20.29 ) |
| 2 | 25.54 ( 22.01 - 28.78 ) | 0.71 ( 0.59 - 0.85 ) | 20.44 ( 17.93 - 24.46 ) |
| 3 | 26.11 ( 22.13 - 29.75 ) | 0.69 ( 0.61 - 0.78 ) | 17.53 ( 15.34 - 20.71 ) |
| 4 | 35.17 ( 30.40 - 39.87 ) | 0.50 ( 0.42 - 0.59 ) | 18.01 ( 15.56 - 21.13 ) |
| 5 | 26.79 ( 20.87 - 32.87 ) | 0.63 ( 0.54 - 0.72 ) | 15.29 ( 12.95 - 18.08 ) |
| 6 | 27.25 ( 20.56 - 34.82 ) | 0.63 ( 0.52 - 0.73 ) | 17.85 ( 15.23 - 21.62 ) |
| 7 | 27.44 ( 21.09 - 34.25 ) | 0.62 ( 0.53 - 0.71 ) | 12.67 ( 10.69 - 15.52 ) |
| 8 | 27.53 ( 18.83 - 35.34 ) | 0.62 ( 0.51 - 0.75 ) | 15.16 ( 12.70 - 18.69 ) |
| 9 | 23.45 ( 15.12 - 30.82 ) | 0.68 ( 0.58 - 0.79 ) | 11.45 ( 9.54 - 14.47 ) |
| 10 | 20.05 ( 9.84 - 31.18 ) | 0.72 ( 0.57 - 0.86 ) | 15.24 ( 12.30 - 19.92 ) |
| Posterior mode and 95% credible interval for the inheritance regression: 0.12 (0.06 - 0.17) |  |  |  |

### Random regression of size

To model the family of size-at-age functions in bighorn sheep ewes, its genetic basis, and associated phenotypic and genetic covariances of size across age, we fitted a random regression animal model (Kirkpatrick et al., 1990, 1994; Meyer & Hill, 1997; Meyer, 1998; Wilson et al., 2005) of the form

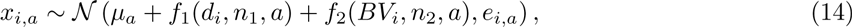

where *x_i, a_* is the size of individual *i* at age *a* and *μ_a_* are age specific intercepts. *f*_1_ and *f*_2_ are random regression functions on natural polynomials of order *n*, for permanent environment effects and additive genetic values, respectively. The permanent environment effect refers to all consistent individuals effects other than the additive genetic effect (see Kruuk & Hadfield, 2007). In both *f*_1_ and *f*_2_, *n* was set to 2, allowing the estimation of random intercepts, slopes, and curvatures. Polynomials were applied to mean-centred and standard deviation-scaled ages to improve convergence. Finally, heterogeneous residuals across ages were estimated (*e_i, a_*). *d* and *BV*, vectors with individual and pedigree values, respectively, were assumed to follow normal distributions, 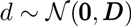 and 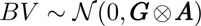. Both 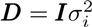, where 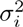 is the permanent environment effect of individual *i*, and the additive genetic variance-covariance matrix, ***G***, are 3 × 3 matrices, ***A*** is the pedigree-derived additive genetic relatedness matrix, and ⊗ denotes a Kronecker product. More information on partitioning of phenotypic variance into different components of variation using general pedigrees and the animal model is provided by Lynch & Walsh (1998), Kruuk (2004) and Wilson et al. (2010). To obtain the genetic variance-covariance matrix for the 10 ages, the following equation is used

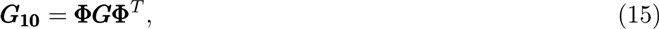

where ***G***_**10**_ is the resulting 10 × 10 genetic matrix, ***G*** is the 3 × 3 genetic matrix estimated by the model and Φ is a 10 × 3 matrix with the polynomials evaluated at each age (Kirkpatrick et al., 1990; Meyer, 1998). A 10 × 10 matrix, ***D***_**10**_, for individual effects at the 10 ages can be obtained similarly. The estimated fixed effects and variance parameters are presented in table 2.

**Table 2:**
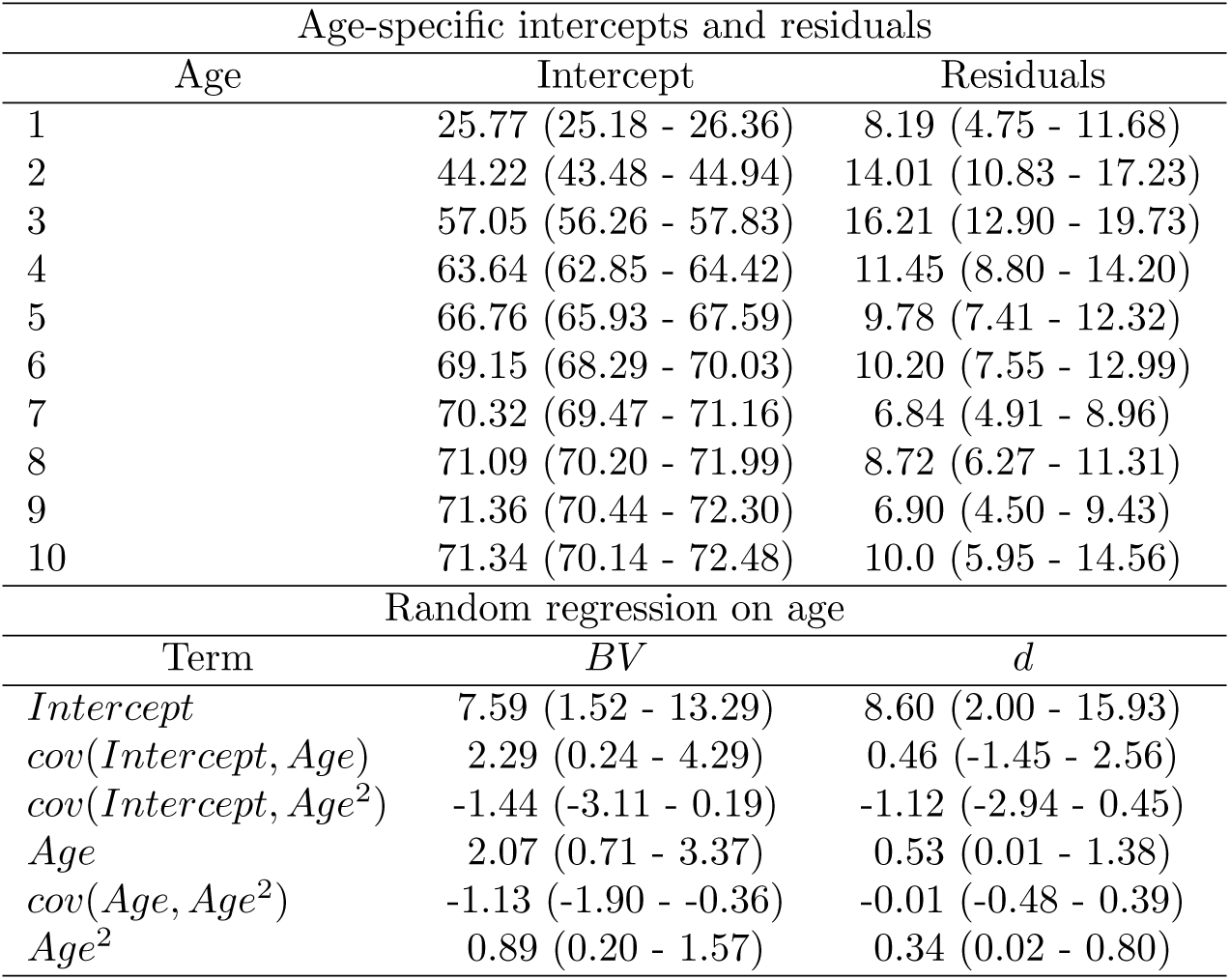
Coefficients for the random regression model on body mass, including age-specific fixed intercepts and residuals (upper part), as well as estimates for the intercept, slope (*Age*) and curvature (*Age*^2^) of the random effects on breeding values (*BV*) and permanent environment (d, lower part) for the bighorn sheep population of Ram Mountain. The values correspond to posterior modes and 95% quantile-based credible intervals.

### Recovering resemblance within and across-generations

We compare the correlations in mass among ages implied by the development functions typically adopted in IPMs and those derived from a RRM, to the observed phenotypic correlations (Figure 8A-C). We used the path rules, as described for the theoretical models, to obtain the correlation matrix for size at different ages implied by the IPM approach. There was no need to do the same for the RRM, as these correlations were recovered with equation (15). We also analyze the proportion of correlation recovered for different gaps between ages (projection steps, Δ𝓉) by both models (Figure 8D). The RRM estimates a phenotypic correlation matrix (Figure 8C) that is much more similar to that observed (Figure 8A) than the correlation matrix implied by the IPM approach (Figure 8B). Across-age correlations are better recovered by the RRM than by the IPM approach (Figure 8D). The proportion of correlation in size among ages recovered by an IPM follows the pattern predicted in figure 2, with high recoveries for a single projection step, and then rapidly decaying to near zero (Figure 8D). As predicted by our theory, typical parameterizations of the development functions severely underestimate similarity of trait values within individuals across ages.

**Figure 8:**
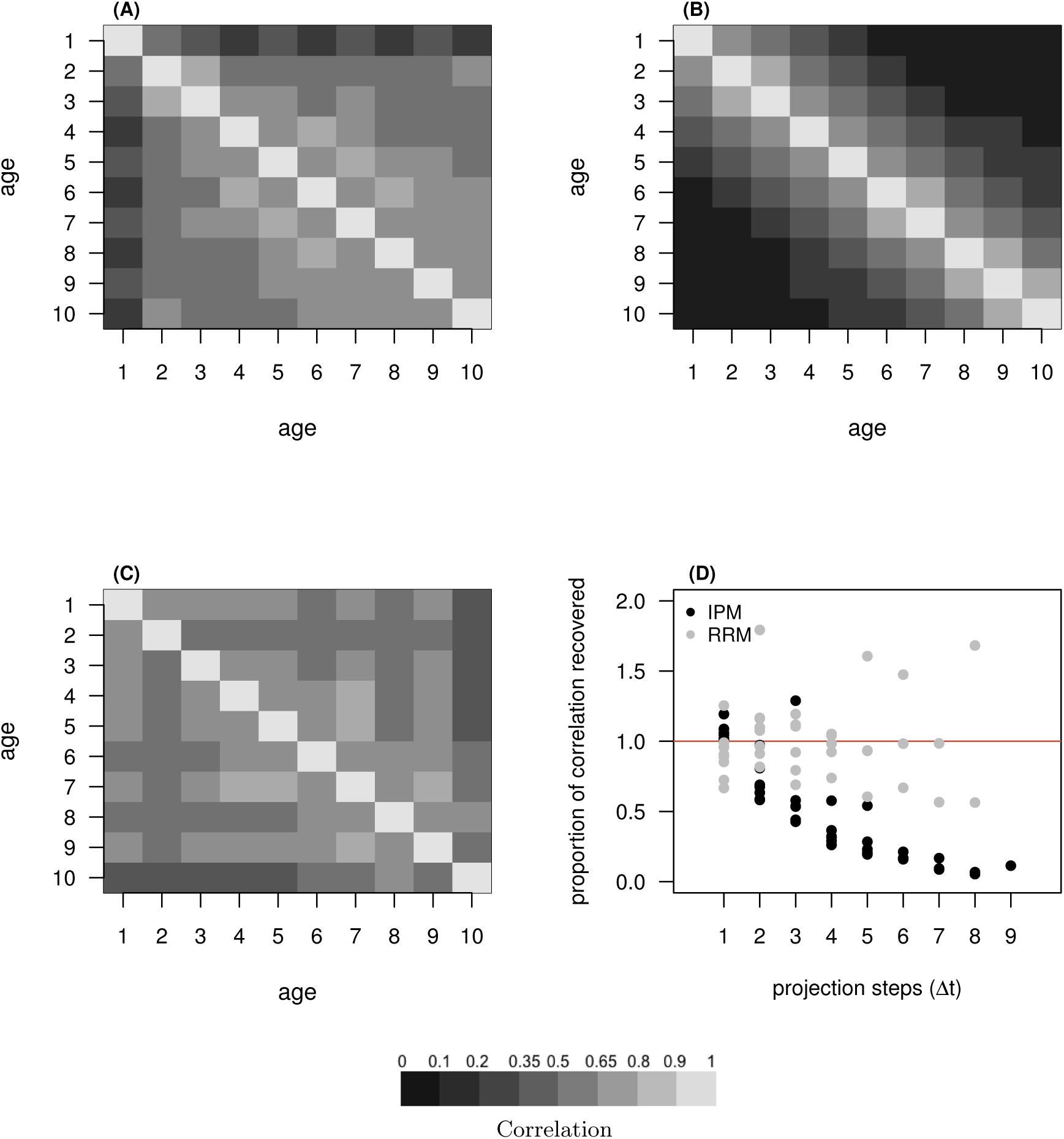
Observed phenotypic correlation matrix for size across ages for the bighorn sheep population of Ram Mountain (**A**), and analogous matrices implied by the IPM (**B**) and estimated by the RRM (**C**) approaches. Proportion of the correlations in size among ages recovered by the IPM (black dots) and RRM (grey dots) for different age gaps (projection steps, Δ𝓉), using the observed phenotypic correlations as reference (**D**). In (**D**), a porportion of 1 (horizontal line) corresponds to a perfect recovery of the observed correlation.

Second, we show the parent-offspring regressions recovered by the RRM and the IPM, and use the “observed” regressions as reference (Figure 9). These latter values correspond to regressions of daughter mass on maternal mass for all matching ages, also including random intercepts for mother ID by age, year and cohort, as well as heterogeneous residuals by age. The cross-age biometric inheritance function implemented in IPMs recovers parent-offspring regression for lambs (age 1), but for older ages most similarity between parents and offspring is missed (Figure 9). In contrast, the patterns of parent-offspring similarity recovered by the RRM are of the observed order of magnitude throughout most of the life cycle (Figure 9).

**Figure 9:**
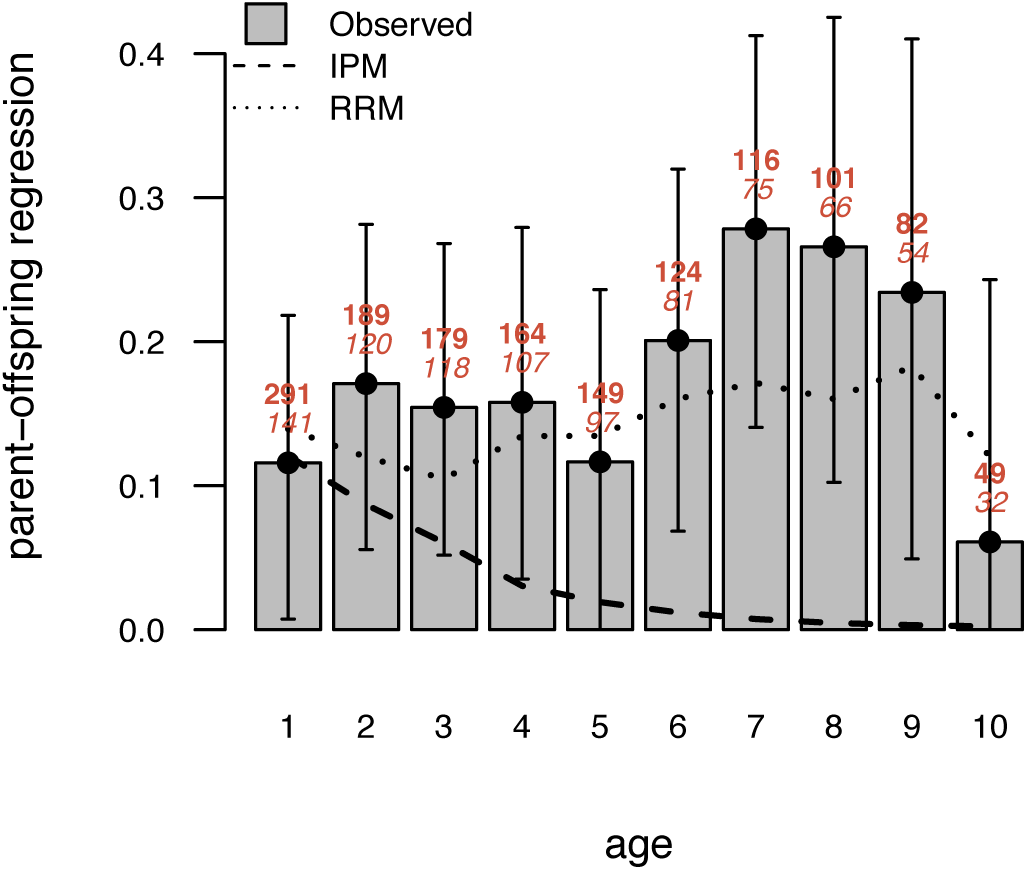
Parent-offspring regressions estimated for different ages for the bighorn sheep population of Ram Mountain, by the IPM and the RRM approaches. The observed values, and the corresponding 95% credible intervals, were estimated by a linear mixed model of daughters’ mass on mothers’ mass for matching ages, with random intercepts for the mother ID by age, year, and cohort. The values on top of the bars correspond to the number of offspring (top, bold) and mothers (bottom, italic) available for each age.

## Discussion

We have shown analytically that IPMs, as typically implemented, will generally, and often severely, underestimate quantities that are critical to evolutionary inference. Both our theoretical results and our empirical example show that phenotypic covariances within and across individuals can be effectively zero in these models, due purely to artifacts of their construction. Additionally, the static nature of the inheritance function (parent-offspring regressions with fixed intercept) artificially reverses any response to selection. Consequentially, IPMs, as typically constructed, will inevitably suggest that evolution is not an important aspect of the dynamics of traits over time. We suggest, and demonstrate empirically, alternative approaches that could be used to characterize some key functions in IPMs. IPMs in principle are extremely useful and highly flexible, and their original conceptualization (Easterling et al., 2000; Ellner & Rees, 2006) should be broadly compatible with a variety of alternative ways of characterizing variation in growth and inheritance.

The main reason why development functions in IPMs fail to recover within-generation covariances of traits is regression to the mean. This problem is well-understood in evolutionary and ecological studies (e.g. Kelly & Price, 2005). In IPMs, this problem is particularly severe because the multiple age-specific projection steps compound the effect of measurement error to reduce covariance among predictor and response variables. Consequently, covariance between non-adjacent ages, which can be substantial (Figure 8A, Wilson et al., 2005), is severely underestimated (Figure 8B), even when measurement error is relatively small (Equations 1 and 2).

The failure of biometric inheritance functions to predict phenotypic similarity among relatives is partially also a direct manifestation of regression to the mean. Indeed, it is the canonical manifestation of regression to the mean - coined in exactly this context by Galton (1886). What we now understand is that Mendelian factors are inherited, and that, in terms of statistical mechanics of quantitative genetics, environmental variation can be regarded as measurement error obscuring the influence of breeding values. Any model of inheritance that does not include our understanding of how inheritance drives similarity among relatives in quantitative traits (Fisher, 1918, 1930; Wright, 1922, 1931) cannot be expected to suffice for even the most basic evolutionary predictions. Another issue arises from assuming that the biometric inheritance function is constant. Whenever the mean phenotype changes, the intercept of the parent-offspring regression necessarily changes as well. To presume that the intercept of the parent-offspring regression is constant across generations constrains the mean phenotype to be able to respond only transiently to selection, as we show by analytically iterating the mean phenotype in a simple IPM model structure (Figure 7D). We reiterate that our criticism of a constant inheritance function is not a criticism of models assuming a constant heritability, whether that heritability is modelled using a genetical (i.e. using constant 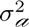 and 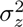) or a biometric approach (i.e. using a parent-offspring regression with a constant slope). Rather, the key point is that the mean phenotype cannot evolve in a model where a parent-offspring regression has a fixed intercept.

In our theoretical models, we use simple but general development and inheritance functions that are specifically designed to isolate these two fundamental processes from each other. In practice, however, the undesirable behaviours that we have modelled separately will interact. Importantly, in iteroparous organisms, where multiple episodes of reproduction occur over the lifetime, regression to the mean in development functions will further obscure relationships between parents and offspring, with increasing effects as parents age (Chevin, 2015). Additionally, biased estimates of covariance of parents and offspring are compounded across multiple generations. The underestimation of similarity between parents and offspring will be compounded at each generation, leading to increasingly severe undervaluation of the relevance of relationships among more distant relatives to the evolutionary process. This interaction is very evident in the empirical example we present. Parent-offspring regressions recovered with the development and inheritance functions generally used in IPMs (Figure 9) could not be predicted by the two-age theoretical model presented here, and specifically by equation (11).

IPMs with typical cross-age biometric inheritance functions have been recommended for studying evolutionary responses to selection (Coulson et al., 2010; Coulson, 2012; Rees et al., 2014). Some studies applying this approach have concluded that non-evolutionary changes in trait distributions are the major contributors to temporal changes in phenotype (Ozgul et al., 2010; Traill et al., 2014). Our theoretical findings do not indicate that these conclusions are wrong. Rather, we demonstrate that these are the conclusions that this kind of model must inevitably generate when applied to any system, regardless of whether evolutionary change is important or not. Since typical parameterizations of IPMs neglect the vast majority of similarity between parents and offspring, they cannot attribute phenotypic change to evolution. Concern about how IPMs model the transmission of dynamic traits had been previously raised (Hedrick et al., 2014; Chevin, 2015; Vindenes & Langangen, 2015; van Benthem et al., 2016). Particularly, Chevin (2015) identified some issues addressed in this paper, presenting insightful numerical examples that illuminate the main concern with the cross-age structure of the inheritance function. Besides our analytical demonstrations, and the numerical examples made available by Chevin (2015), we also provide an empirical example, using random regression analysis to address the issues presented here. The random regression model provided substantial improvement in recovering both correlations across ages within a generation (Figure 8D), and parent-offspring regressions reflecting how breeding values are transmitted over generations (Figure 9).

Vindenes & Langangen (2015) discuss joint models of static traits (constant through life) and dynamic traits (such as those typically handled in IPMs) in the general IPM framework. They suggest that incorporation of static traits could solve some of the problems that had begun to be acknowledged about evolutionary inference with IPMs (Hedrick et al., 2014; Chevin, 2015). The authors propose that the static trait, birth mass in their example, could be modelled as influencing mass at all other ages and demographic rates, which would allow covariances among birth mass and older ages to be well recovered. In a sense, using random regression animal models as we suggest treats breeding values (as opposed to some realized phenotypic value) as a static trait, but critically also models the inheritance of breeding values, not as some observed function, but according to the principles of quantitative genetics. It is noteworthy to mention that a genetic notion of trait transmission has already been implemented into an IPM for a single Mendelian locus (Coulson et al., 2011). The authors constructed an IPM that describes the dynamics of body mass and a biallelic gene determining coat color in wolves (*Canis lupus*). In contrast to biometric IPMs of quantitative traits, Coulson et al. (2011) conclude that the genetic variance within the study population is enough for natural selection to cause evolution. In fact, it is in principle relatively straightforward to implement an IPM that uses the basic principles of inheritance of polygenic quantitative traits to define inheritance functions of breeding values; such exercises have indeed begun for a single trait (Childs et al., 2016). It is easy to conceive of multivariate extensions of such inheritance functions (based on multivariate versions of equations 3 and 4), whereby one could treat age-specific sizes as different characters, and estimate genetic variances and covariances from data. Nonetheless, a great deal of work is still required. For long-lived organisms, genetic covariance matrices of age-specific traits would be very challenging to estimate with useful precision (Wilson et al., 2010). Furthermore, the dimensionality of resulting phenotypes would overwhelm typical strategies for implementing IPMs (Coulson et al., 2010; Rees et al., 2014; Merow et al., 2014). In practice, a key challenge, but a surmountable one, will be to develop sufficiently flexible, low-dimensional characterizations of the genetic basis of development for practical estimation and subsequent modelling. The function-valued trait approach we adopted with our random regression model of bighorn sheep ewe mass is just one such possibility. Other approaches could possibly be even more useful; for example, uses of various autocorrelation functions (Pletcher & Geyer, 1999; Hadfield et al., 2013), or factor-analytic mixed model (de los Campos & Gianola, 2007; Meyer, 2009; Walling et al., 2014).

## Summary

We have shown analytically and using and empirical example that standard implementations of integral projection models will generally severely underestimate the likelihood of evolutionary change. IPMs to date have been constructed using characterizations of development and inheritance that would not stand up to scrutiny in studies focusing on development and inheritance. It is not surprising that more complex models built on such functions behave poorly. In fact, insofar as the ability of IPMs to track the full joint distribution of phenotype has been suggested as their main quality for ecological inference, the problems that preclude their typical use for evolutionary inference should be of equal concern to ecologists. Importantly, we have suggested ways in which more nuanced models of development, and a modern understanding of inheritance, can be incorporated into the general IPM approach. A great deal more work is required before IPMs based on adequate models of development and inheritance will be field-ready. As a next step, careful studies of the performance of different approaches for characterizing the genetic basis of developmental trajectories, with particular focus on approaches that could be incorporated into an IPM framework, are needed.

## Acknowledgments

We thank Jarrod Hadfield, Loeske Kruuk, Luis-Miguel Chevin, Josephine Pemberton, Graeme Ruxton, Jean-Michel Gaillard, and Sandra Hamel for valuable comments and discussions. We are also very grateful to all those who worked on the bighorn program over decades. The bighorn research is supported by the Government of Alberta, the Université de Sherbrooke and an Alberta Conservation Association Challenge Grant in Biodiversity, NSERC Discovery Grants to D. Coltman, M. Festa-Bianchet and F. Pelletier and the Canada Research Chair in Evolutionary Demography and Conservation. M. B. Morrissey is supported by a University Research Fellowship from the Royal Society (London). M. J. Janeiro is supported by a PhD scholarship (SFRH/BD/96078/2013) funded by the Fundação para a Ciência e Tecnologia (FCT).

